# Evolving cohesion metrics of a research network on rare diseases: a longitudinal study over 14 years

**DOI:** 10.1101/030809

**Authors:** Carlos B. Amat, François Perruchas

## Abstract

Research collaboration is necessary, rewarding, and beneficial. Cohesion between team members is related to their collective efficiency. To assess collaboration processes and their eventual outcomes, agencies need innovative methods - and social network approaches are emerging as a useful analytical tool. We identified the research output and citation data of a network of 61 research groups formally engaged in publishing rare disease research between 2000 and 2013. We drew the collaboration networks for each year and computed the global and local measures throughout the period, relating connectivity with the normalized impact. Although global network measures remained steady over the whole period, the local and subgroup metrics revealed a growing cohesion between the teams. Transitivity and density showed little or no variation throughout the period. In contrast the following points indicated an evolution towards greater network cohesion: the emergence of a giant component (which grew from just 30% to reach 85% of groups); the decreasing number of communities (following a tripling in the average number of members); the growing number of fully connected subgroups; and increasing average strength. Normalized citation measures, although ranking the member groups as above average impact for Spanish biomedical output, do not support correlation between collaboration and citation impact. The Spanish research network on rare diseases has evolved towards a growing cohesion - as revealed by local and subgroup metrics following social network analysis. This greater cohesion does not translate into a greater impact, as measured by the citation analysis of the papers.

## Introduction and background

Scientific collaboration has been defined as “interaction (…) among two or more scientists that facilitates the sharing of meaning and completion of tasks with respect to a mutually shared, superordinate goal” (Sonnenwald, 2007). From a purely scientific perspective, collaboration is necessary to cope with the increasingly challenging, ambitious, and demanding objectives of many research initiatives (in terms of human knowledge as well as material resources). Evidence for growing collaboration comes not only from quantitative data on the rising number of individual contributors per paper (Wuchty, Jones and Uzzi 2007) but also from the increasing presence of collective authors. The ATLAS Collaboration in Particle Physics is an extreme example (Cho, 2011). In the biomedical research fields, the MONICA Project [cardiovascular health] of the WHO, or the EPSILON study on schizophrenia are good examples of the collective involvement of different countries, centers, and teams in pursuing common research problems.

Joint research gives competitive advantage to research partners in the form of more frequent citations for co-authored papers (Larivière et al. 2015) and therefore greater funding opportunities. Beyond the strictly science boundaries, collaborative research also has beneficial socioeconomic outcomes and so national and supranational public entities are encouraging research partnership in the implementation of specific programs or by introducing criteria to foster collaborative approaches in funding applications. The cross-national COST actions that “fund pan-European, bottom-up networks of scientists and researchers across all science and technology fields” (http://www.cost.eu/about_cost/how_cost_works), or the Clinical and Translational Science Awards program of the US National Center for Advancing Translational Sciences (https://www.ctsacentral.org/) are good examples of the former; while the principles for funding multi-institutional collaboration in innovation and research published by the UK Research Council (http://www.rcuk.ac.uk/funding/principles/) is a good example of the latter. Fostering agencies need instruments to assess the efficiency of research collaborations. In fact, a recent report on so-called “team science” recommends that researchers “partner with team science leaders to evaluate and improve analytical methods and tools for team assembly” (Cooke and Hilton 2015) and network analysis has emerged both as an adequate framework and as a convenient analytical tool to understand these processes and evaluate their outcomes (Hunt, Whipple and McGowan 2012; Bian et al. 2014).

### Social network analysis of collaboration

A network is a representation of a system. It consists of vertices that represent the entities of the system. Pairs of vertices are joined by edges that represent a particular kind of interconnection between these entities (Estrada 2011). Citation relationships between documents are considered the source of a type of knowledge network, namely, citation networks (Newman 2003b); while the co-authorship of articles in learned journals (Newman 2001b; Newman 2001a) along with the networks constructed from collaborative research grant applications (Bian et al. 2014) have been studied as examples of social networks. Throughout this paper, we refer indistinctly to network (net) or graph; likewise, we do not differentiate between teams and groups that are connected or linked by edges or ties.

In co-authorship networks, individuals or groups (entities) are connected if they have co-authored one or more papers. Such a simple relationship and the resulting systems have attracted a considerable number of works from Scientometrics, Social Network Analysis (SNA), and other research fields. Co-authorship networks have been approached either in a longitudinal or a cross-sectional manner and analyses have been performed at the collective or elemental level. We will next review some examples of these approaches and their utility for our purposes.

Mark Newman can be credited for undertaking the first large-scale analysis of scientific co-authorship (Newman 2001b; Newman 2001a). Although his main aim was to obtain a reliable social network based on the assumption that joint authorship reflects genuine professional interaction between scientists, the metrics he used for characterizing the networks (we will review these in the appropriate section) have remained as a model for analyzing co-authorship networks at the collective level. His study, however, is cross sectional, and reduced to a five-year period with accumulated figures. María Bordons and colleagues, in another example of transversal research, relate the research performance and the network position of individual researchers in pharmacology, nanoscience, and statistics (Bordons et al. 2015). There are two other evaluative papers focused on the participation of research groups in the Clinical and Translational Science Awards: the already mentioned article on the University of Arkansas for Medical Sciences (Bian et al. 2014) uses SNA for “evaluating the impact of resource allocation to different programs”; and an article on the Indiana Clinical and Translational Sciences Institute (Hunt et al. 2012) “derives a single consented ranking of important or influential nodes in a collaboration network”.

Xiaoming Liu and co-workers analyzed the structure of collaboration within the Digital Libraries research community and provided quantitative metrics for the concepts of status and influence of individual authors (Liu et al. 2015). They added to the individual prestige metrics some general measures of the whole structure of the network; however, their analysis remains static. A good example of a longitudinal study at the elemental level could be the analysis of “social inertia” in which Ramasco and Morris (2014) follow 14 research collaboration networks (and another derived from the Internet Movie Database) during an indefinite time period.

Shortly after the seminal work by Newman, several other statistical physicists studied the evolution of social networks using co-authorship networks as examples (Barabási et al. 2002), although to our knowledge Newman’s paper was the first example of dynamic analysis of co-authorship, their main interest seemed to be the large scale modeling of complex evolving networks. In contrast, the work by Katy Börner and others (2005) appears to be the first that combines the positional metrics of individuals in the network with several “success” indicators of the papers they contribute and, more importantly, a longitudinal follow-up of the characteristics of the whole network. In the same line, Luis Bettencourt and co-authors follow the emergence and development of a series of research specialties tracking several co-authorship network metrics as the corresponding fields evolve (Bettencourt, Kaiser and Kaur 2009). More recently, Ghosh and collaborators followed several structural measures of evolving co-authorship networks (Ghosh, Kshitij and Kadyan 2014) however; their treatment of network cohesion is quite superficial.

Several sets of metrics have been applied, in the above mentioned and other works, to characterize either the behavior of the whole co-authorship network and its evolution, or the role of individual nodes whose topology reveals an outstanding position, influence, or importance in the observed research field. These network and node metrics have been combined with “efficiency” estimates, usually in the form of research outputs and impact indicators that attempt to confirm the benefits of collaboration. However, before we move on with a detailed review of these measures and state our objectives, it is worthwhile describing the context in which our analysis takes place.

### CIBERER as our case of study

Biomedical Research Networking Centers (CIBER after the Spanish acronym) is a Spanish public initiative to support single-topic research on specific broadly-defined disease or health problems. Following an initial call in 2006, nine monographic centers were established on Neurodegenerative Diseases, Hepatic, and digestive diseases, Public health and epidemiology, Bio-engineering, biomaterials and nanomedicine, Diabetes and associated metabolic disorders, Physiopathology of obesity and nutrition, Mental health, Respiratory diseases and Rare diseases. This last mentioned center, CIBERER for short, is the object of our work.

The word “center” might be misleading, as every consortium is made up of a number of research teams from various parent organizations. From its beginning in 2007, 61 research teams joined CIBERER and, after one was removed in 2009, the consortium included 60 teams, encompassing 700 people in 2013. The last annual report details an annual budget (in 2013) of 6.7 million euros coming from the hosting public institutions and research program funding plus 1.5 million euros from private funding, which comes to a total of 8.2 (or almost USD 9.4 million at the current exchange rate).

The starting point for the CIBER program is “the need to boost research excellence through the implementation of stable structures of research collaboration” (Ministerio de Sanidad y Consumo 2006). This main goal adds to a specific feature of rare diseases that is implicit in its name: this is a very large group of diseases (over 7,000 following international criteria) with a low prevalence in the population. Most are oncological, neurological, or metabolic disorders – usually with genetic origins. So it should come as no surprise that the need for networked research on rare diseases had already been stressed (Aymé and Schmidtke 2007). Collaboration in rare diseases research comes, then, as a twofold necessity. Firstly, as the number of patients is small, it is necessary to cooperate to avoid the fragmentation of research and gain shared knowledge. Secondly, it is necessary to cooperate to associate the clinical features and, eventually, identify new genes or associate gene expression and mutation consequences to the clinical features of a given disorder. Hence, the need for organizational plus instrumental collaboration adds to the call for “excellence” through the foundation and development of institutional collaboration. How could we measure both the degree of collaboration and the effects eventually resulting from joint research efforts? The next paragraph introduces some concepts from network analysis and metrics that may help represent collaboration relationships and measure their strength and evolution. For the sake of comprehension, the concepts used are loosely-defined.

Collaboration and co-authorship are far from equivalent concepts (Laudel 2002) although Scientometric practice has sanctioned the use of the latter as a proxy for the former. Here, we refer to the relationship between two investigators who jointly appear on the byline of a research paper as co-authors. When aggregated to the institutional or even higher level, as is the case with this article, we refer to collaboration as, say, two universities collaborating rather than co-authors collaborating in one or more papers. Our network takes teams as the entities of analysis (nodes or vertices in the SNA terminology) and establishes connections (edges) between pairs of teams if their members have co-authored one or more papers. Co-authorship edges show some intensity according to the number of coauthored papers between teams. The total number of papers co-authored by a team is its strength or weighted degree. As an example, let us look at the small network on the left side of Figure 1. The vertex labelled ‘b’ connects with three other vertices (a, c and d) in the way that vertex d does; but b has more strength despite having the same degree because the weight of the edge b-a is two. Both have degree 3 but b has strength 4. Although the strength distribution gives some idea about the relationship among entities, a better understanding of the whole network can be drawn from group level metrics – especially if these metrics are followed along the same period. On the right side, Figure 1 depicts the same net but at a more advanced stage and several changes are quite evident. The most obvious is that the two components on the left (vertices h and i) have coalesced into one component that connects all the vertices. Another less obvious change is the appearance of triads: sets of three fully connected vertices from a previous situation where they were connected only in couples. This, for example, is the case of vertices d-e-f and f-g-d. Finally, it is obvious that a complete subnet has been developed among the four vertices d to g, with each connected to the others.

**Figure 1.**
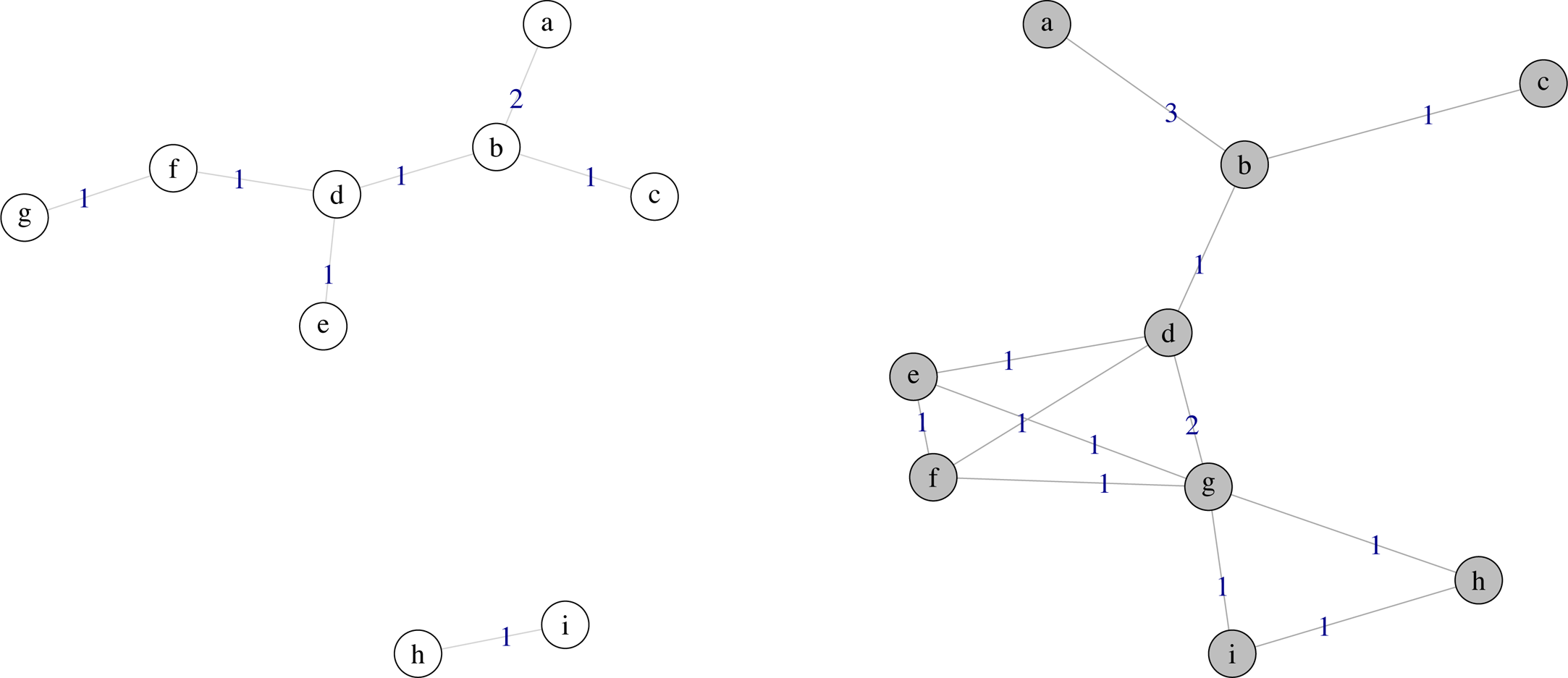
Successive stages of an ideal network evolving (from left to right) towards a greater cohesion.

Network performance is related to network cohesion. Several reviews and metaanalyses on cohesion-performance relationship have found a positive correlation, although this correlation is moderate and highly dependent on intragroup processes (Chiocchio and Essiembre 2009). We approach the concept of group cohesion in a loose and pragmatic way, as the inclination to forge, maintain, and even reinforce social ties in order to achieve common goals and mutual benefits. With respect to research performance, we follow common practice and use citation impact as a proxy, notwithstanding our awareness of the inherent limitations of this approach.

A network is a network insofar as it connects its members. The main purpose of this paper is to follow the collaboration relationships among the research teams of a formal research consortium on rare diseases and examine if the collective evolution leads to greater cohesion. This will be accomplished by first determining the research output of the groups and identifying the journal publications that two or more groups co-author. Secondly, we will build the collaboration networks and apply network metrics to observe the collective evolution of the groups and their interrelationships. We are particularly interested in those measures that best reflect the above definition of network cohesion. Thirdly, we will obtain some performance measures, namely, normalized citation frequencies of the papers, in order to explore some eventual association between network cohesiveness and the citation impact of research publications. Finally, we will offer some concluding remarks which could orientate future research on scientific co-authorship networks.

## Dataset and analysis

We studied networks established among CIBERER research teams that co-authored papers. Although CIBERER was formally established late in 2006 and started activities in 2007, we extended the scope of our study from 2000 because it was important to identify some “invisible college” effect, that is, some previous interactions already in place among the member groups. In that case, the foundation of collaboration could be hardly attributable to the formal network activities.

Using author and affiliation data, we identified and retrieved from the Web of Science (WoS) 4710 journal publications contributed by CIBERER teams during the period 2000 to 2013. As the resulting bibliographic lists used to be incorporated to the web site of CIBERER (www.ciberer.es) on a periodical basis, every group had the opportunity to chek for the accuracy and completeness of their own bibliographies and to ask eventually for amendments.

In 987 of these papers we identified two or more co-authoring CIBERER teams, meaning that the authors of these papers were affiliated to two or more groups. As we were only interested in the collaboration inside this consortium, we disregarded any other co-authoring data in our analysis, although the impact analysis will take into account the presence of international contributors.

After processing the bibliographic dataset, we obtained two tabular files for every year in the period 2000-2013. The first contained the groups who published at least one paper in that year. Along with their identification, this vertex file listed every team along with their specific research areas, the geographical location of their host institutions, their clinical or basic orientation, and the number of papers contributed. Only teams publishing at least one paper in a given year were included in the corresponding set. The second file was a plain list of the pairs identified in the papers for the same year. Those familiar with network analysis would recognize the two components of the standard file format used by most applications. We used the R package iGraph (Kolaczyk and Csardi 2014) to run the analytical routines on the data, which included some very basic distributions, such as the degree distribution, as well as network cohesion metrics as they evolved in the time frame.

In determining the impact of co-authored papers, Börner chose a fractional approach to calculate the impact of every author and distributed the number of citations received by a paper among its co-authors (Börner et al. 2005). We focused, however, on the papers and made a whole count approach because we wanted to observe if co-authored papers have a greater impact than single team papers. We also obtained from the WoS, the citation data of every Spanish paper published between 2000 and 2011. This data included not just the citation frequency but also the chronology of received citations. The impact indicator we used is a variant of the so-called item oriented field normalized citation score (Lundberg 2007) which compares the relative number of citations to publications from a specific group to the average citations received by the Spanish papers of the same document type and subject area that were published in the same year. In addition to using national instead of world publications as the reference set, we limited the citation window to the year the paper was published plus the two following years. Thus, a paper in a journal in the category Neurosciences (say PMID 17070050) received no citations in 2007, the year it was published, and accumulated 6 + 7 = 13 in the following two years. Spanish articles published in 2007 in the same subject category were cited on average 7.53 times in the same period and so the quotient results in 13/7.53 = 1.73, meaning that this particular paper had a citation impact 73% above average (which is 1).

Research performance is affected by the internationalization of research – meaning the contribution of foreign research teams to the projects and papers (Inzelt, Schubert and Schubert 2008). To control for this effect, we isolated those papers with no foreign contribution and compared the scores of papers coauthored by several groups with those contributed by a single team in search of some significant differences using the Mann-Whitney test. We also analyzed the distribution of citation scores for the collaborative and single group papers during the period.

## Results

### Research output

Before discussing the network analysis, it seems convenient to give some data on the research output of CIBERER and its groups. We identified 4710 journal publications contributed by the 61 research teams between 2000 and 2013. The Web of Science subject categories most frequently attributed to the papers were by far Genetics & Heredity and Biochemistry & Molecular Biology, then came Endocrinology & Metabolism, Neurosciences, Clinical Neurology, and Oncology. This distribution gives some idea of the cross-disciplinary composition of the consortium.

Figure 2 summarizes the research output of the groups in terms of published papers per team during the period. The average number of papers per team and year grew from 4.46 to 10.1 over the period with standard deviations of 2.97 and 9.46, respectively; these figures are consistent with the evolution of the median – of the distribution: in 2000 half of the groups published four papers at least – while in 2013 they doubled their output and published eight papers. Dispersion grew progressively during the period; the initial interquartile range was four, while leaving aside the years 2008 and 2009, the range reached and even exceeded seven by the final year of the series. Another imbalance is revealed by the progressive appearance of outliers who, in the two latter years of the period, reached a maximum output of more than 50 papers a year. It seems that, regarding their publications output, setting up the consortium caused a growing inequality among the groups.

**Figure 2.**
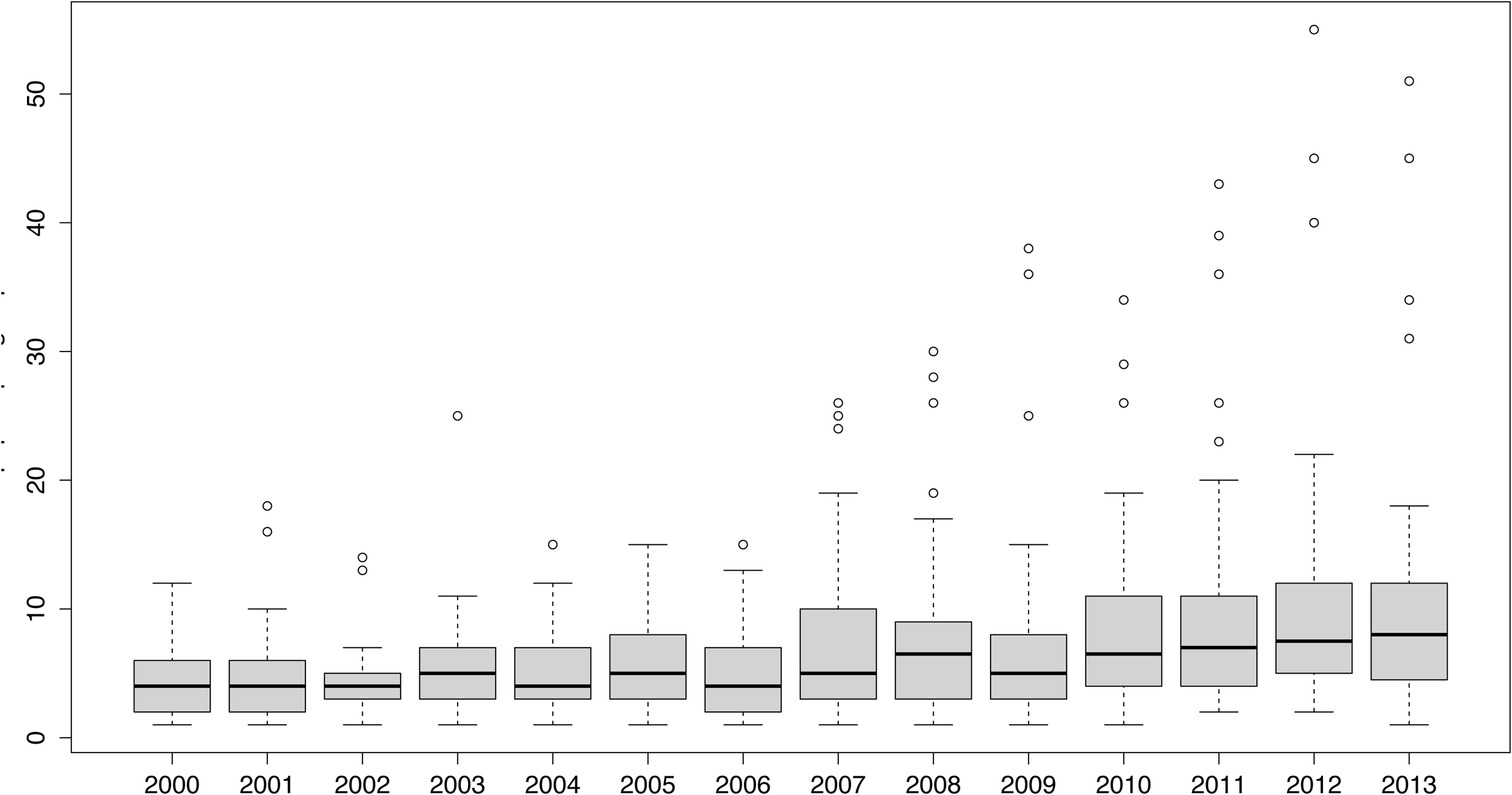
The research output of CIBERER in terms of the distribution of papers per group for each year of the period. Full data is available from the authors.

On the other hand, Figure 3 presents the dramatic increase in the number of foreign institutions (here expressed on a logarithmic scale) contributing to CIBERER papers in the last years of the period. For 2000 to 2002, half of CIBERER papers just showed one foreing collaborating institution (log10 = 0). The median of the distribution rose to 0.3 (meaning two foreing institutions) between 2003 and 2011 while, at the same time, the outliers proliferated. In the last two years of the series, half of the papers had three international contributing institutions or more and the great number of cases which outlay have displaced the arithmetic mean above four foreign institutions per paper. The extreme case in 2012 corresponds to a consensus paper (PMID 22966490) with more than 1,150 contributing institutions.

**Figure 3.**
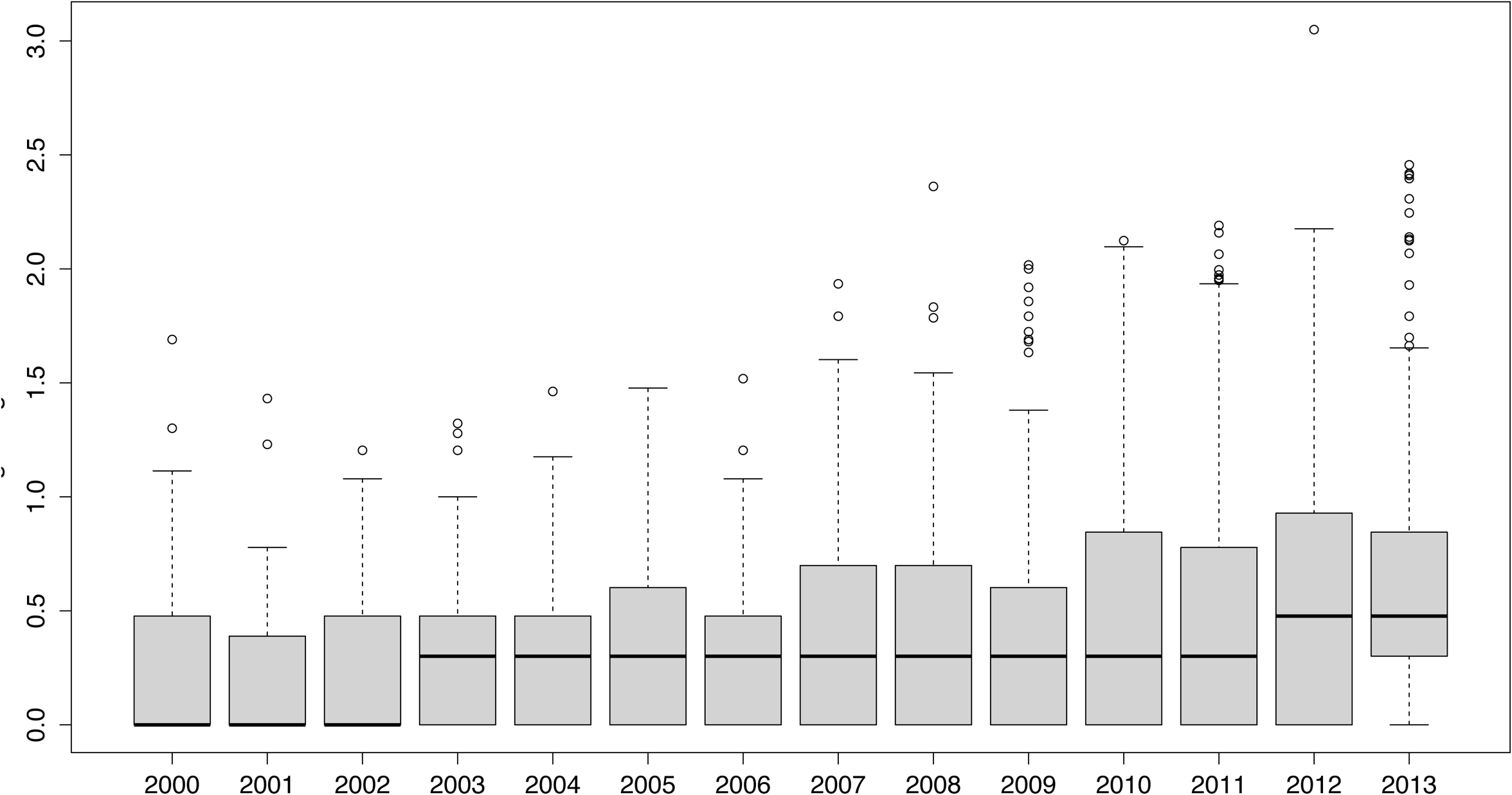
The frequency distribution of the number of foreign institutions per paper on a logarithmic scale shows its dramatic increase, particularly in the two last years. As always, full data is available upon request.

### Cohesion analysis

Many co-authorship network analysesis begin by looking over the number of connections incident to the nodes. While this approach is reasonable in cross-sectional studies focusing on individual elements of the net, centrality metrics looking to identify outstanding nodes are of little use in studying the collective behavior of evolving networks. It makes much more sense to pay attention to cohesion metrics because those values reflect the evolution of CIBERER groups towards either coalescence or disaggregation. Table 1 gives a summary of the activity of the teams and also of their relationships throughout the period., Three series of values are provided for every year. The first one includes global measures, the second one relates to the local features of the evolving network; finally, a set of organizing patterns is detected in the shape adopted by some topological features. The first row in Table 1 contains the number of active teams for each year, meaning teams that published at least a one paper in that year. Since the formal constitution of the consortium in 2007, almost the 97% of the groups are active. The share was much lower in 2000, with less than 86%, and rose to a 93.4 % in 2006. By 2007, the year the shared activity took off, all the groups where active in publishing research papers and this feature has remained stable except for 2011 and 2013.

**Table 1.**
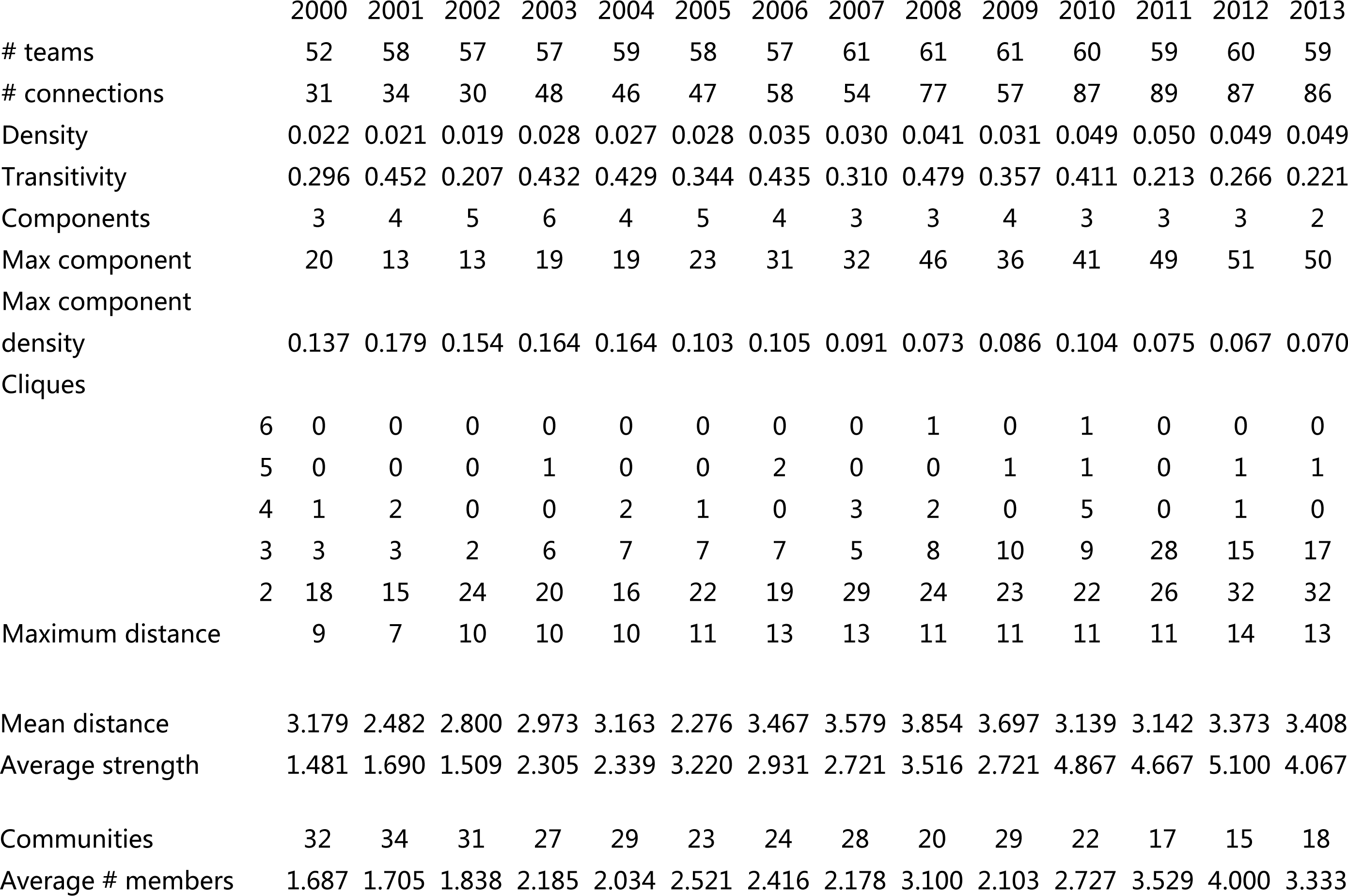
Global, local and community metrics for each annual co-authorship network

### Global measures

Although most studies use the number of co-authoring authors or groups as the main indicator of the intensity of collaboration, it is the number of connections that best reveals how a network evolves. In fact, the average number of groups per paper was 1.13 in 2000, 1.15 in 2007, and 1.18 in 2013 (confidence interval 1.13-1.21 for this last year) while the number of connections almost tripled for the number of active groups which grew 10 % between the extreme years. Indeed, the average number of co-authored papers, denoted by the average strength row, grew from less than three in 2007 to more than four in 2013 after peaking in 2012 (while the period 2000-2006 presents more modest figures). Figure 4 shows the evolution of the distribution of the tie strength among the groups.

**Figure 4.**
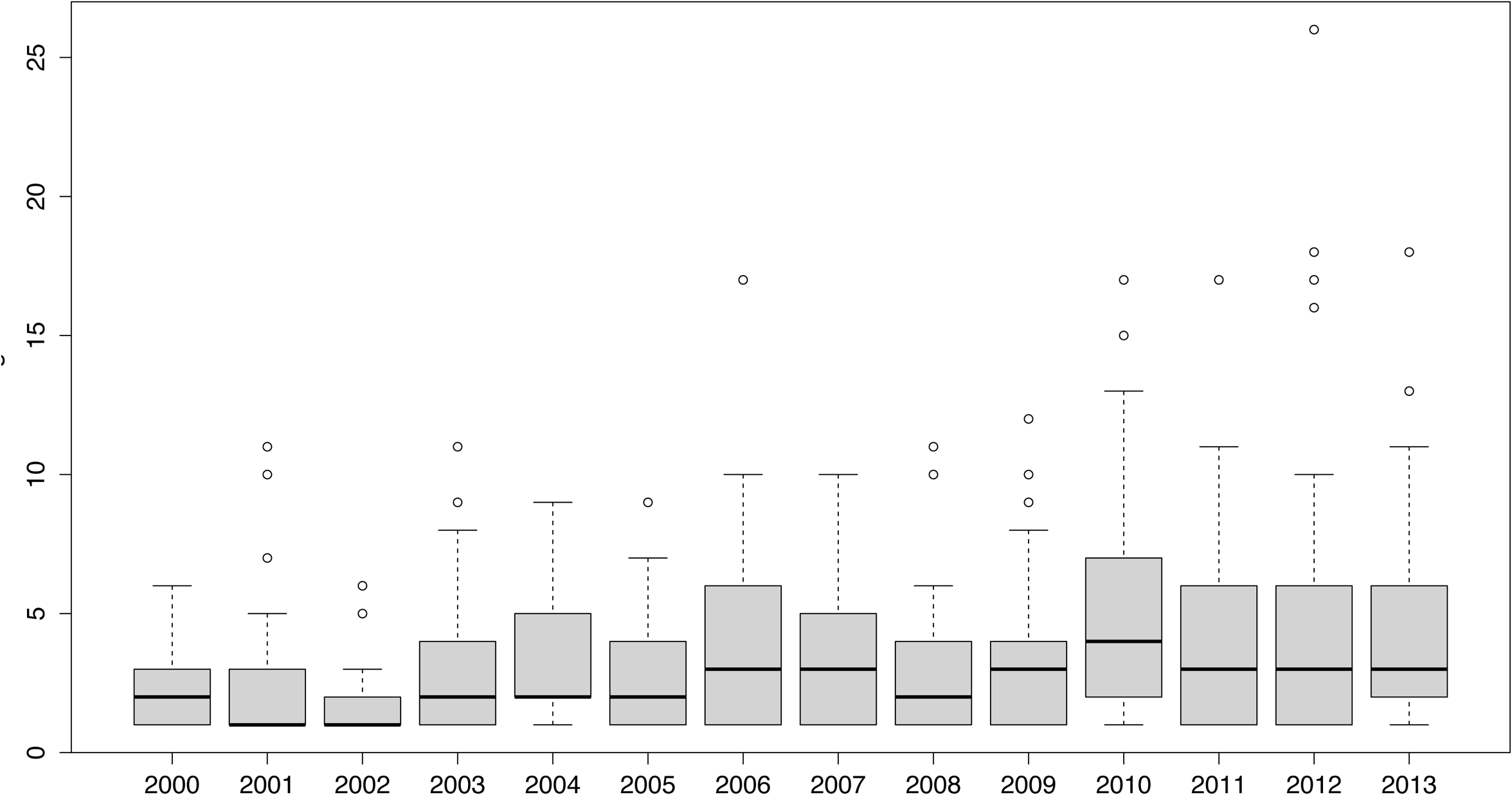
The evolution of vertex strength (weighted degree) of the groups along the period. Full distribution is available from the authors.

The proportion of connections established to the total number of possible connections is called the network density and, in our case, this density increased 2.23 times during the period – however, it multiplied by 1.59 between 2000 and 2006 and by a mere 1.63 in the formal period. A commonly used quantity in network literature is the ratio of the total number of triples that form triangles (as with the g-h-i nodes in Figure 1 right) over the total number of connected triples, as with a-b-c). This is termed transitivity or clustering coefficient, and the values fluctuate during the period with no clear tendency.

### Local measures and subgraphs

A network is connected if there is a path (a sequence of nodes and ties) between each pair of nodes. Figure 1 (right) is a connected network while the network on the left has two components. It is obvious that the more components, the less cohesive is a network. The giant component is the subgraph with the larger number of connected groups. The corresponding values for these two metrics are shown in Table 1 and although the number of components has remained practically unchanged since 2007, it is the size of the giant component that reveals how cohesive the CIBERER network has become. In 2007, the first year of collective activity, the giant component involved 52.46 % of the active groups; in 2013 this share rose to 83.33%. In the early years, only 2006 showed a share above 50%. The emergence of a giant component is not the only indicator at the local level of a rise in connectivity and, hence, net cohesion. The graph is complete when every node is connected to every other node. A complete subgraph is called a clique, as is the case with subgraphs d-e-f-g and g-h-i in Figure 1 (right). Cliques in Table 1 are classified by the number of interconnected teams – from two to six. It becomes apparent that the formal period is populated by cliques of three to five groups, with cliques of order six in 2008 and 2010.

### Communities and mixing

In addition to the local topology of a network and its subgraphs, there are some results related to the patterns of interrelation among the nodes, the so-called communities. A community is formed by an internally connected set of nodes for which the internal density is significantly larger than the external density (Estrada 2011). The last two rows of Table 1 contain data on the evolving community structure of the CIBERER network. The most obvious finding is the sharp decline in the number of communities since the consortium’s start and an increase in the average number of members. This result is consistent with the increase in higher order ( ≥ 3 members) cliques and the average number of members of each community. To further investigate this logical organization of the net, we examined whether there was a tendency for the groups to connect with alike others. This feature is called homophilly (or assortativity) and, like the correlation coefficient, its values range from −1 to 1, depending on the degree of association of nodes with respect to some nominal or scalar variable (Newman, 2003a). We have calculated the assortativity coefficient of the networks for 2000, 2007 and 2013 with respect to the degree of the vertices and several attributes such as output (number of papers), location, research field, and basic or clinical profile. With regard to this last feature, teams were classified according to their host institutions. So, a group pertaining to a research institution (say Spanish High Research Council) or to an academic one (Autonomous University of Madrid, for instance) were classified as basic gropus while those working in hospitals were considered as clinically oriented.

The values appear in Table 2 and show (both with respect to the number of collaborations of every group and the number of published papers) values close to zero – meaning a lack of influence on the propensity to collaborate. Most of the research groups were concentrated in the three cities included in the table and it is worth noticing that the coefficients captured in 2000 decrease in 2007 and 2013. Geographical proximity does not seem to play a significant role after an initial stage in the first period. A similar decline in the coefficients can be seen with regard to the clinical or basic orientation of the groups but, in this case, negative values might suggest a propensity to collaborate between basic and clinical groups, that fits with the objectives of the consortium. Moreover, the adscription of some groups to the same research line does not translate into a greater connectivity; as the values captured at the extreme years decrease. The most conspicuous variation appears in the case of hereditary cancer, a highly assortative set of seven groups in 2000 whose thematic relationship seems to have vanished as time went by. This is also the case for the pediatric groups; however, it started from a lower assortativity coefficient based on its thematic profile.

**Table 2.** Assortativity coefficient with regard to geography, subject field and profile of the CIBERER research groups

|  | 2000 | 2007 | 2013 |
| --- | --- | --- | --- |
| assortativity (degree) | 0.035 | 0.034 | -0,029 |
| research output | -0.093 | -0.008 | 0.066 |
| geographical location |  |  |  |
| Barcelona | 0.32 | 0.41 | 0.291 |
| Madrid | 0.354 | 0.366 | 0.181 |
| Valencia | 0.347 | 0.232 | 0.155 |
| clinical or basic profile | -0.408 | 0.175 | -0.031 |
| subject field |  |  |  |
| Mitochondrial medicine | -0.071 | 0.388 | 0.166 |
| Genetic medicine | 0.077 | 0.112 | 0.175 |
| Hereditary Metabolic Medicine | 0.262 | 0.51 | 0.101 |
| Neurosensory pathology | 0.139 | 0.232 | 0.051 |
| Pediatric and Developmental |  |  |  |
| medicine | 0.466 | 0.25 | 0.078 |
| Hereditary cancer | 0.65 | 0.19 | 0.27 |
| Endocrine medicine | NA | -0.29 | -0.06 |

### First dates

When we observed the year in which two groups first co-authored a paper, we found that more than a half (54.67%) of these 300 “first acquaintances” took place in 2007 or later, which reveals the influence of the consortium on the behavior of the groups.

The three-part Figure 5 captures three moments of the evolving collaboration among CIBERER groups. Research groups are represented as circles whose diameter is proportional to their research output. As only the collaborating groups are depicted, it is evident from these layouts the progressive incorporation of the units to the commong goal and the growing connectivity from earlier stages.

**Figure 5.**
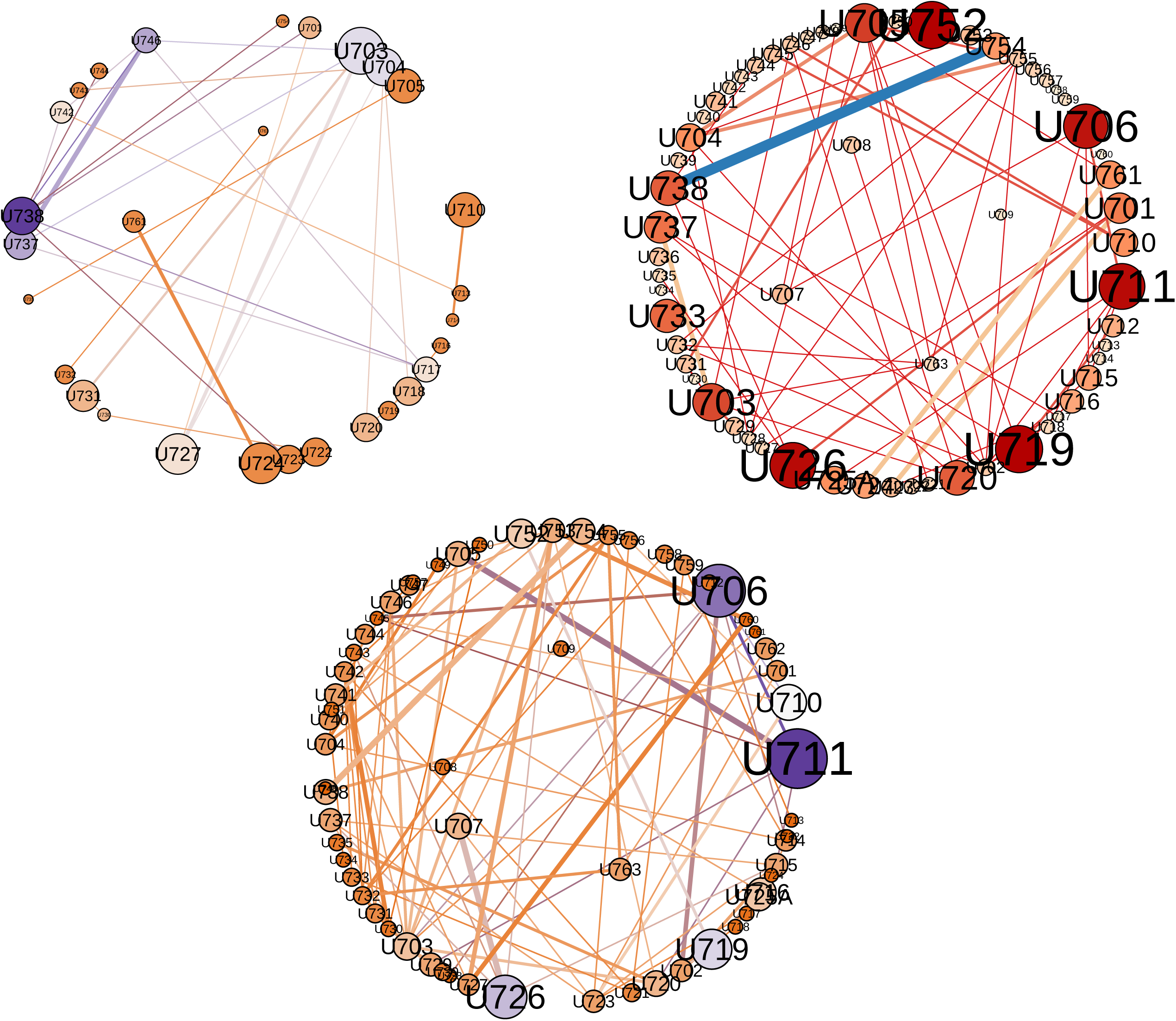
Three snapshots of the CIBERER co-authorship network. Clockwise from top left, the nets from 2000, 2007 and 2013. In these dual circle layouts (drawn with Gephi) nodes represent teams and have been sized following the research output of every group. Isolated (non collaborating) teams have been removed. Thickness of edges denotes intensity of collaboration. Nodes can be identified by their labels.

### Citation impact

Except for 2003, CIBERER papers performed quite well in terms of weighted citation frequency as mesaured through the Karolinska indicator (Lundberg 2007) here calculated for a three-year window. The mean values outperform the average of the corresponding subject fields particularly in 2002 (Ki=1.449) and in 2008 (Ki=1.424). However, as Figure 6 points out, the median values never reach the unit (marked as the horizontal dotted line) meaning that half of the papers don’t get the expected citation score. This fact, along with the coincidence of heavy ouliers in those succesful years, suggests that, as expected, the citation distributions are highly skewed. For example, another consensus document contributed by 236 institutions and published in 2008 (PMID 18188003) had 405 citations in the three-year window after publication. When compared to Figure 3, the citation distributions of Figure 6 suggest a finding that several groups (Larivière et al. 2015 for example) have confirmed: the relationship between coauthorship and impact data.

**Figure 6.**
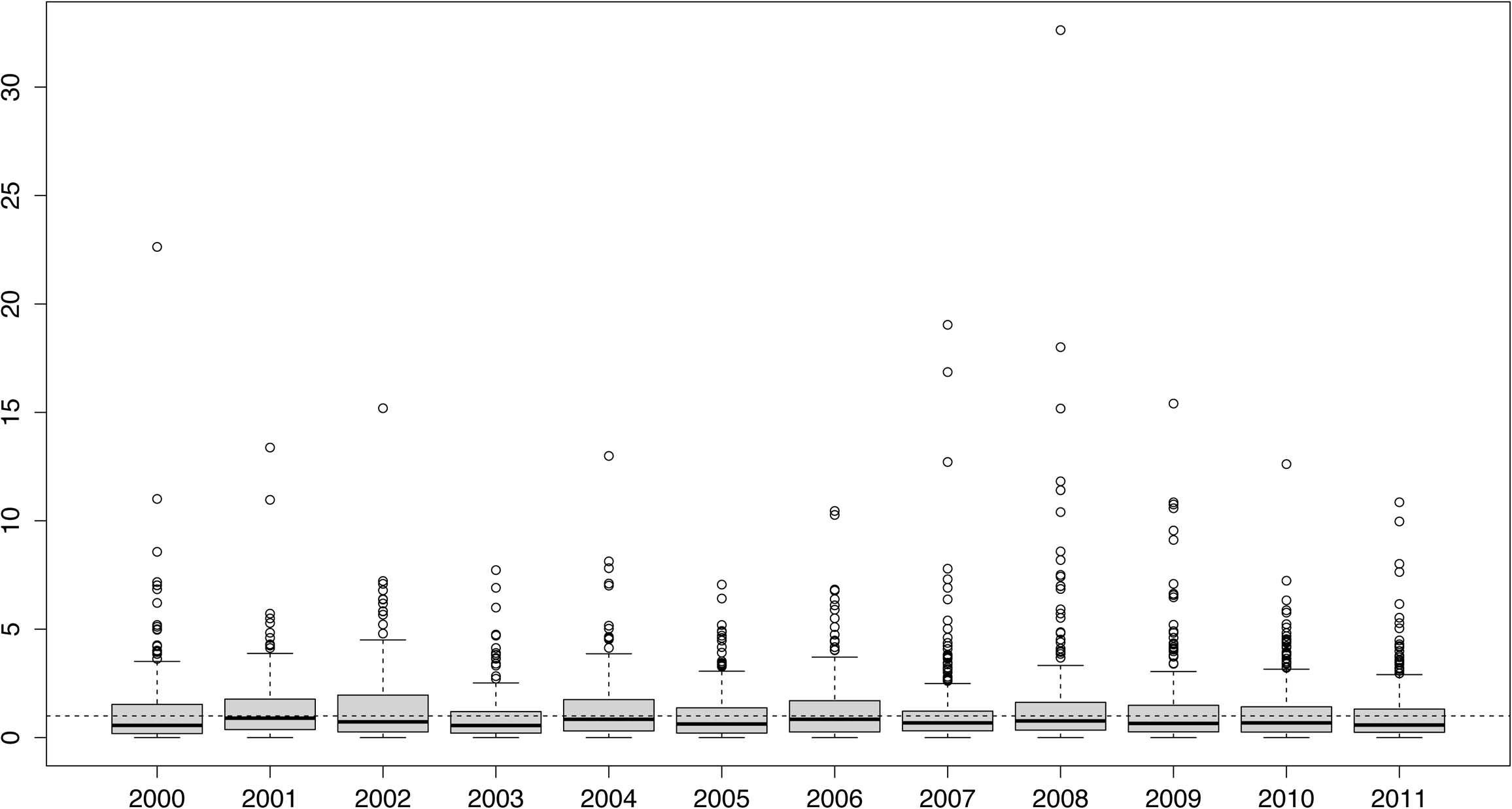
Evolving impact metrics of CIBERER papers from 2000 to 2011. The scores have been calculated following Karolinska indicator (Lundberg 2007) and the doted horizontal line marks the average impact of the Spanish papers in the corresponding subjects fields.

To further investigate this matter and control for the effect of international collaboration on citation impact, we focused our citation analysis on the papers published by CIBERER groups between 2000 and 2011 and not co-authored by foreign institutions. Out of the 2,340 papers which met these conditions, 328 were contributed by more than one CIBERER group while 2,012 were authored by a single team. Let us call these collaborative and singleton papers respectively. Rather than using the mere citation frequency, we drew again on the field normalized citation impact of each paper (Karolinska indicator or Ki for short) as formulated in the methods section (Lundberg 2007).

The top left density plot in Figure 7 depicts the frequency distribution of Ki for the 2,340 papers. Three years after publication, 325 papers (approximately 14%) remained uncited while a similar proportion (13.77%) were cited just once. Nine papers were cited 50 or more times in the three-year period that followed their publication. This data comes as no surprise as citation distributions are usually skewed. For the sake of comparison, the overall proportion of uncited Spanish biomedical papers (those attributed to any of the 74 Web of Science biomedical subject categories) in the same period was 19.79%.

**Figure 7.**
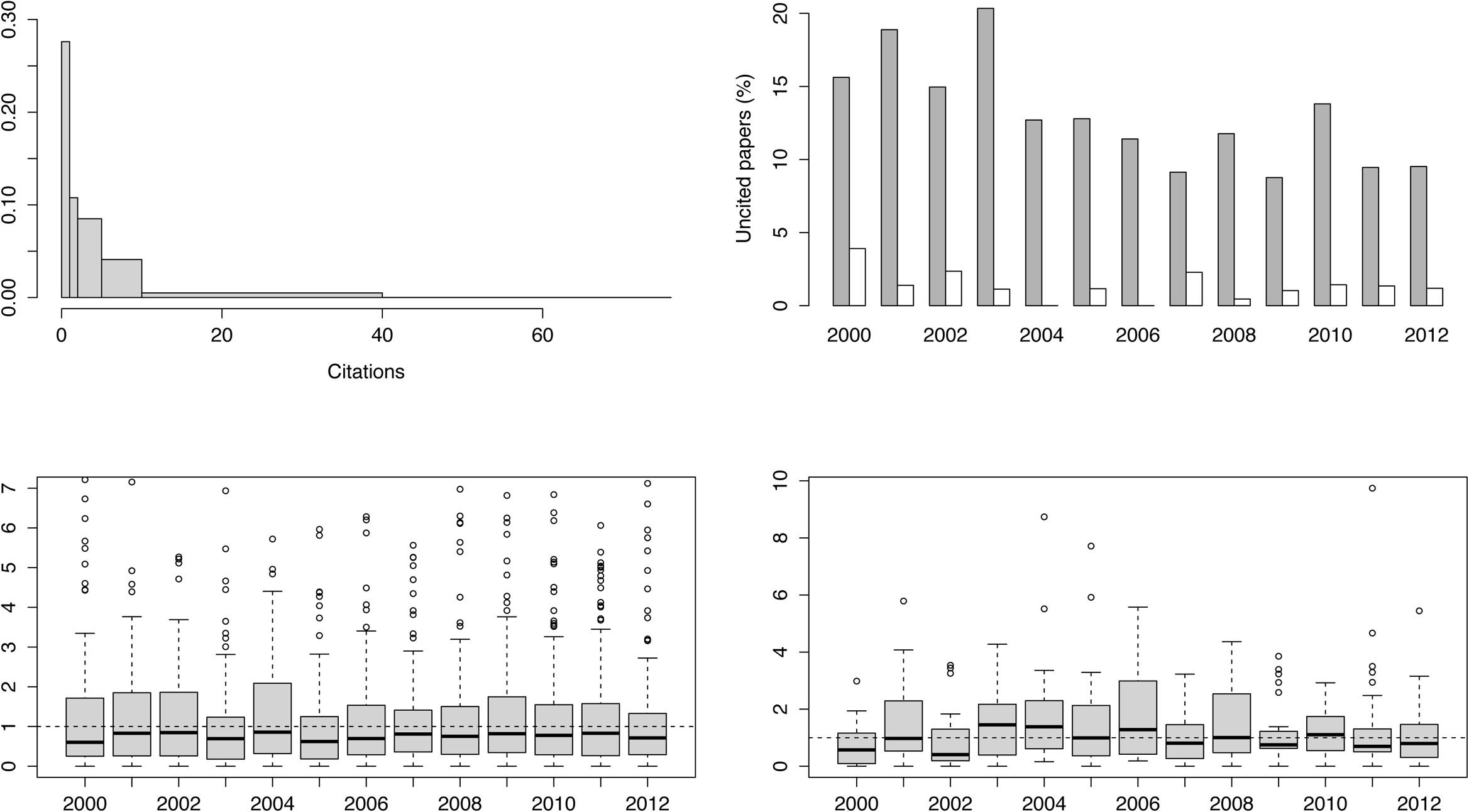
A summary of the citation impact analysis of CIBERER. Clockwise from top left the density plot of the citation frequency of the papers; the share of uncited papers among the singleton (dark bars) and the collaborative papers; the distribution of the Karolinska index (Ki) from 2000 to 2012 of singeton papers (bottom left) and that of the collaborative papers (bottom right). Full data are available from the authors.

To compare the citation impact of both collaborative and singleton sets, our first approach was to observe the proportion of uncited papers over those published every year in each of the two groups (Figure 7, top right). On average, the proportion of single group papers uncited in the three-year term was 10.41 % for those published in 2007 and onwards – while the share rises to 13% if we take the whole period from 2000. The percentages for the collaborative group of papers were 1.29 and 1.15 respectively. Therefore, the share of uncited papers was approximately ten times lower among the collaborative papers.

The Wilcoxon test for the two independent groups (the Mann-Whitney test) revealed significant differences between the Ki distributions for the collaborative and singleton sets of papers (W = 330680, at p-value = 0.01108) with an index (mean ± standard deviation) of 1.3 ± 1.4 and 1.2 ± 1.51 respectively. The collaborative group exceeded the singles by 1.51 to 1.21 in the first period. In the latter period, the two groups showed similar values (1.2 to 1.18 for the collaborative papers).

A closer view of the citation distributions is given in the two boxplots at the bottom of Figure 7, which show how the Ki of the papers is distributed throughout the initial and formal periods of CIBERER. The bottom left plot corresponds to the singleton papers and the bottom right to the collaborative group. The dotted horizontal line through both plots marks the unit value, or the level where Ki equals the average citation rate of the subject categories attributed to the papers. The first difference among the two distributions is the reduced number of outliers for the collaborative papers. There seems to be a tendency towards coalescence and homogeneity in this group while the singleton shows a greater spread of Ki values. With regard to the singleton distributions between the two periods, 39.2 % of the papers published in the initial (2000-2006) period had a Ki greater or equal to the unit. The share was similar (40.55 %) in the latter period. By contrast 54.42 % of the collaborative papers published in the initial period had a Ki larger than one and this falls to 40.88% in the formal period of the consortium.

These last results do not suggest that collaboration among CIBERER research groups does not translate into greater impact. It must be taken into account that the papers with no foreign contribution represent barely half of the research output. Indeed, collaboration is just one of the factors that determine citation impact. Moreover, the initial call for groups already required a certain impact from previous publications. Finally, the taxonomy of network or group performance is much richer than a pure measure of academic influence could reflect (Chiocchio and Essiembre 2009) and so, the citation impact might be related just to the contextual performance while not influencing the task and outcome performances (which are closer to the final goals of the network).

### Conclusions

Two main conclusions arise from this study. Firstly, the analysis of Spanish research network on rare diseases reveals a growing cohesion. Secondly, this greater cohesion does not translate into a greater impact, as measured by the citation analysis of the papers.

We have detected some level of interaction among the groups before the net was formally founded. However, and despite the apparent steadiness that some global measures indicate, there is a clear tendency towards coalescence as revealed by the progressive reduction of the number of separate components along with the emergence of a giant component. Indeed, the appearance of fully connected subgroups (cliques) of higher order and the reduction of the number of separate communities combined with the increase in the average number of members are additional indicators of greater network cohesion. In the end, a network is made up of connected elements, otherwise it is not a network.

The normalized citation score of the papers reveals a larger citation impact of CIBERER groups over the rest of contemporary Spanish publications in the same fields. This advantage was present before the formal foundation of the net and is not affected by the participation of more than one group when the contribution of international teams is disregarded.

Finally, let us quote the team science report that says “As teams and groups develop and move through their phases of scientific problem-solving, their interactions will change, and the field must identify how to measure these team processes”. Bibliometrics and co-authorship network analyses are explicitly mentioned, along with other qualitative methods such as the techniques whose combination is needed to ascertain how team processes are related to the multiple goals of transdisciplinary team science (Cooke and Hilton 2015 p. 52) Our ultimate goal has been to contribute with our results to this emerging field of research.

## Aknowledgements

The Spanish Ministry of Economics and Competitiveness partially supported this research (grant number ECO2014-59381-R).

